# ShadowCaster: compositional methods under the shadow of phylogenetic models for the detection of horizontal gene transfer events in prokaryotes

**DOI:** 10.1101/684704

**Authors:** Daniela Sánchez-Soto, Guillermin Agüero-Chapin, Vinicio Armijos-Jaramillo, Yunierkis Perez-Castillo, Eduardo Tejera, Agostinho Antunes, Aminael Sánchez-Rodríguez

## Abstract

Horizontal gene transfer (HGT) plays an important role in the evolution of many organisms, especially in prokaryotes where commonly occurs. Microbial communities can improve survival due to the evolutionary innovations induced by HGT events. Thus, several computational approaches have arisen to identify such events in recipient genomes. However, this has been proven to be a complex task due to the generation of a great number of false positives and the prediction disagreement among the existing methods. Phylogenetic reconstruction methods turned out to be the most reliable but they are not extensible to all genes/species and are computationally demanding when dealing with large datasets. On the other hand, the so-called surrogate methods that use heuristic solutions either based on nucleotide composition patterns or phyletic distribution of BLAST hits can be applied easily to genomic scale, however, they fail in identifying common HGT events. Here, we present ShadowCaster, a hybrid approach that sequentially combines compositional features under the shadow of phylogenetic models independent of tree reconstruction to improve the detection of HTG events in prokaryotes. ShadowCaster predicted successfully close and distant HTG events in both artificial and bacterial genomes. It detected HGT related to heavy metal resistance in the genome of *Rhodanobacter denitrificans* with higher accuracy than the most popular state-of-the-art computational approaches. ShadowCaster’s predictions showed the highest agreement among those obtained with other assayed methods. ShadowCaster is released as an open-source software under the GPLv3 license. Source code is hosted at https://github.com/dani2s/ShadowCaster and documentation at https://shadowcaster.readthedocs.io/en/latest/.

## Introduction

Lateral or horizontal gene transfer (HGT) plays an important role in the genome evolution and ecological innovation of prokaryotic communities. HGT in bacteria and archaea occurs more frequently between closely related species than in distant lineages [1]. Transferred genes usually provide selective advantages for survival in the recipient lineage and are kept for long periods of time [2]. Rates of HGT events involving genes critical for survival, growth, and reproduction are particularly high among members of microbial communities that need a quick adaptation to complex environments such as contaminated soil or water [3]. Detecting HGT has been a major focus of attention to better understand microbial evolution. However, it has been proven to be a complex and challenging task.

HGT detection tools can be divided into two main classes: parametric and phylogenetic methods. Parametric methods search for sections of a putative recipient genome that greatly differ from the mean nucleotide composition based on metrics such as oligonucleotide frequencies, GC content or codon usage. Genes exhibiting highly compositional fluctuations from the mean are called atypical/alien genes and their origin is expected to be exogenous. Phylogenetic methods integrate information from multiple genomes and find evolutionary incongruencies while reconciling gene trees with the reference species tree [4]. Both parametric and phylogenetic methods are based on the hypothesis that acquired genes bring “perturbations” to the recipient genome, and that such perturbing signal can be differentiated from the background noise (fluctuations in the recipient genome that arise from other scenarios either than the HGT events), regardless the age of the event.

Up to date, there is no unique method capable of detecting all the HGT events of different ages in a recipient genome because each method has their own advantages and limitations. Phylogenetic methods are best at identifying ancient HGT events provided a large number of orthologs and the generation of a reliable species tree [5, 6]. Evolutionary processes other than HGT, *i.e.* gene duplication and differential gene loss, can also explain incongruences between genes and species trees, which hinder the performance of phylogenetic methods [7, 8]. Parametric methods deal best with recent HGT events that result in noticeable perturbations to the recipient genome mean nucleotidic signature. Over time, however, gene amelioration dilutes the signal of HGT events due to the quick lost of nucleotide compositional differences which drastically reduces the detection power of parametric methods [9]. In presence of gene amelioration parametric method’s predictions become inaccurate and should be validated by phylogenetic approaches, in order to reduce both false positive and negative predictions [10, 11]. Nevertheless, explicit phylogenetic methods (tree building-based) are not practical at large-scale (*e.g.* comprising entire gene repertoires), because they are computationally expensive and time consuming [12]. In this sense, several methods having rather implicit phylogenetic approaches, have arisen. These methods are mainly based on phyletic distribution derived from BLAST searches in order to speed up alien gene detection in multiple genomes/taxa since they bypassed gene tree reconstruction e.g. DarkHorse [13], HGTector [14] and HGT-Finder [12].

Table 1 shows the state-of-the-art of HGT detection methods that rely on different information sources and when applied to the same dataset often generate partially overlapping predictions. Discrepancies among tools might be resolved by combining parametric and phylogenetic methods. It is still unclear what could be the best way to combine different methods without increasing the false discovery rate (FDR) [4, 15].

**Table 1.** State-of-the-art of computational approaches for horizontal gene transfer detection with emphasis in prokaryotic genomes.

| Classification | Implementation | Methodological highlights | Application Domain | Reference |
| --- | --- | --- | --- | --- |
| *Parametric methods* |  |  |  |  |
| **Nucleotide composition** | Alien Hunter<br>(http://www.sanger.ac.uk/science/tools/alien-hunter) | Uses Interpolated Variable Order Motifs (IVOMs) coupled to a Hidden Markov Model (HMM) to detect “alien” genes. | bacterial genomes | (Vernikos and Parkhill, 2006) |
| | No implementation available | Detects “alien” genes based on k-mer ($k = 8$) frequencies using a one-class support vector machine (SVM). | viral, archaeal and bacterial genomes | (Tsirigos and Rigoutsos, 2005) |
|  | No implementation available | Combines two compositional features using a Kullback–Leibler divergence metric to improve the detection of “alien” genes. | artificial genomes | (Becq *et al.*, 2010) |
|  | GOHTAM<br>(http://gohtam.rpbs.univ-paris-diderot.fr/) | Uses a Jensen-Shannon divergence metric from window or gene-based signature data to detect “alien” genes. | prokaryotic and eukaryotic genomes | (Ménigaud *et al.*, 2012) |
|  | No implementation available | Detects “alien” genes based on the selection of nine compositional features using a SVM. | bacterial genomes | (Xiong *et al.*, 2012) |
| **Nucleotide composition plus information from the genomic context** | No implementation available | Implements a multiple-threshold approach to detect “alien” genes from compositional features and genomic context information to reduce the chance of misclassification. | artificial genomes | (Azad and Lawrence, 2011) |
| *Implicit phylogenetic methods* |  |  |  |  |
| **Phyletic distributions based on BLAST searches** | DarkHorse<br>(http://darkhorse.ucsd.edu/) | Calculates a lineage probability index from BLAST searches to predict “alien” genes. | prokaryotic and eukaryotic genomes. | (Podell and Gaasterland, 2007) |
|  | HGTFinder<br>(http://cys.bios.niu.edu/HGTFinder/<br>HGTFinder.tar.gz) | Calculates a horizontal transfer index from BLAST searches to predict “alien” genes. | prokaryotic and eukaryotic genomes. | (Nguyen *et al.*, 2015) |
|  | HGTector<br>(https://github.com/DittmarLab/HGTector) | Establishes statistical thresholds to detect genes that do not adhere to *a priori* defined hierarchical evolutionary categories inferred from BLAST searches. | artificial, prokaryotic and eukaryotic genomes. | (Zhu *et al.*, 2014) |
| **Hybrid methods** |  |  |  |  |
| **Nucleotide composition complemented with an implicit phylogenetic model** | ShadowCaster | See further | prokaryotic genomes | This work |

Here we developed an open-source software called ShadowCaster consisting in the sequential incorporation of parametric and phylogenetic information to improve the detection of HTG events in prokaryotes. By combining two compositional features with an implicit phylogenetic model that defines species relatedness from the number of shared orthologs, ShadowCaster predicts HGT with higher accuracy that the most popular state-of-the-art computational approaches. The core of our method is a likelihood function inferred by phylogenetic shadowing, is used to predict how likely is that an ‘alien’ gene had been vertically inherited. We applied ShadowCaster to predict genome-wide close and distant HGT events in artificial and bacterial genomes. The hybrid nature of ShadowCaster allowed predicting new HGT events in real world data demonstrating its improved performance relatively to pure parametric and phyletic methods. In addition, ShadowCaster predictions showed the highest agreement with those obtained by other methods.

## Design and Implementation

ShadowCaster is an open-source software capable of detecting HGT events in prokaryotes by performing three major tasks: (*i*) ‘alien’ genes identification, (*ii*) phylogenetic shadowing model construction, and (*iii*) per gene likelihood calculation expressing how likely an ‘alien’ gene has been vertically inherited. There are two main components in ShadowCaster to carry out these tasks: one parametric and one phylogenetic. The parametric component uses the genome of the query species to extract nucleotidic compositional information during task (*i*), while the phylogenetic component starts from the proteomes of the query and of other related species covering a diverse spectrum of phylogenetic distances, to perform tasks (*ii*) and (*iii*).

### Parametric component

For the identification of alien genes, we implemented a parametric component that combines two features, each using a particular metric to measure the compositional difference between each gene and the entire genome. Features were implemented as suggested by Becq et al. [6]. The first feature corresponded to the gene length normalized tetranucleotide (4-mers) frequencies with Chi-square as metric and the second was the codon usage with Kullback-Leibler as metric. The per-gene values of each feature were compared to the corresponding value calculated from the entire coding sequence of the genome (*i.e.* a concatenation of all genes in the species).

To classify each gene as “alien” based on features values, One-class support vector machine (SVM) was used since this method is able to perform the outlier detection in an unsupervised fashion. The goal of the One-class SVM (Fig 1B) was to separate the native genes (*i.e.* those with a composition that does not differ greatly from the one of the entire genome) (gray dots in Fig 1B), from atypical genes (green and red dots in Fig 1B) by the estimation of a support distribution. This model requires the selection of a kernel function (radial basis function in our case), and the specification of the bounds of the fraction of support vectors and training errors to use as defined by the user via the ‘nu parameter’ (see technical details in [21] and references therein). In our implementation, the nu parameter can be tuned by the user depending on the type of HGT events to detect, *i.e.* close, medium-far and far events (see below how the nu parameter affects the predictions made by ShadowCaster). Genes classified as outliers by the One-class SVM represent the group of “alien” genes that constitute the output of the parametric component and one of the inputs of the phylogenetic component.

**Fig 1.**
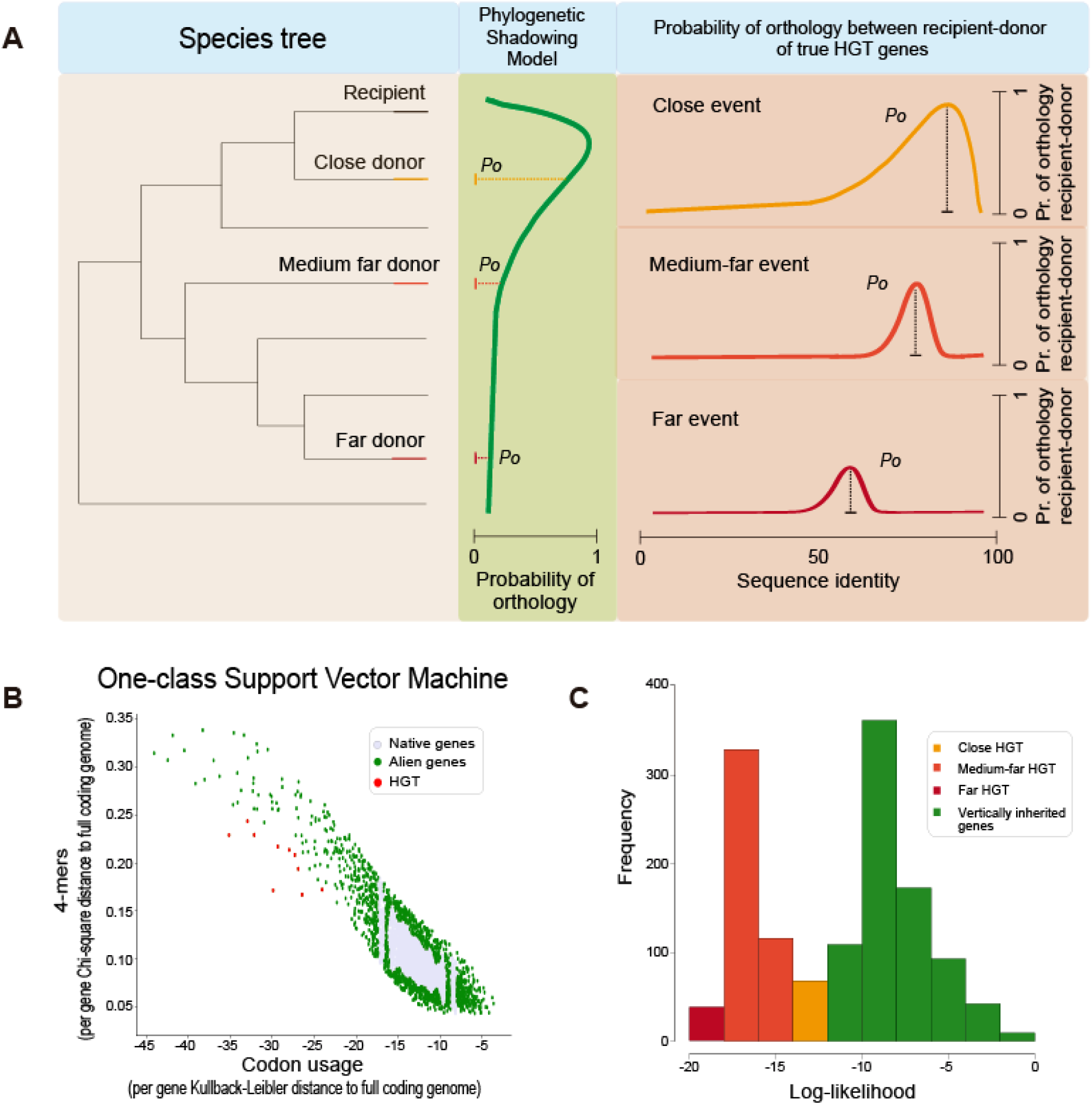
Graphical representation of the conceptual model behind ShadowCaster and of its main outputs. **A:** Probability of true orthology between three gene pairs shared by a recipient species and three phylogenetically related donor species, respectively (species tree in the left-hand side). True orthology probability values (*P*_*0*_) are sampled from probability distributions according to: vertical inheritance (phylogenetic shadowing model defined by the number of orthologs sharing recipient-donor species at different phylogenetic distances, green panel); and lateral inheritance (HGT model, gradient colour curves from orange to deep red represent the *P*_*0*_ distribution in true HGT events occurring at different phylogenetic distance along the species tree. *P*_*0*_ decreases in both vertical and lateral inheritances with the increase of the phylogenetic distance but this behavior is markedly less in HGT. **B:** The distribution/separation of typical (in grey color) and atypical (in red and green colour) genes achieved by the parametric component of ShadowCaster (4-mer frequency and codon usage). **C:** Log-likelihoods for all alien genes detected by the parametric component in a given recipient genome. Log-likelihoods and *P*_*0*_ are related by the equation 2.

### Phylogenetic component

The list of ‘alien’ genes detected by the parametric component is still putative and might contain spurious results (e.g. native genes that did not arise from true HGT events). The aim of the phylogenetic component is then to create a filter capable of differentiating genes that truly arose from HGT events from those native genes that show an atypical composition but that were vertically inherited. The model behind ShadowCaster (Fig.1A) builds on the previous knowledge that the amount of true orthologous genes shared between a given species pair decreases with the phylogenetic distance that separates them. In such scenario, a vertically inherited gene in a recipient species, will show a decreasing homology with its orthologs in increasingly distant species. If one sees the homology between two proteins as a proxy of orthology, and constructs a probability function across a species phylogenetic tree will see a shadowing shape as represented in Fig 1A (green curve). A horizontally inherited gene will in contrast disrupt the shadowing shape of the orthology probability function creating a peak around the acceptor species (see the orange, light red and deep red curves in Fig 1A).

We took advantage of the contrasting behavior of vertically versus horizontally inherited genes in terms of orthology probability distribution across the species tree. While, these two events share similar values of orthology probability *P*_*0*_ (top parts of the green and orange curves, Fig 1A) in close-related species; as far the homology between donor and acceptor species in the vertical inheritance (green curve) get increased, *P*_*0*_ gradually decreases. By contrast, *P*_*0*_ decrease is not markedly evident in medium and far HGT events (light red and red curves, Fig 1A). With such considerations, then we applied a Bayesian inference to estimate how likely is that a given gene has been vertically inherited.

When applied this to our phylogenetic shadowing model, Bayes’ theorem for probability distributions can be expressed as follows:

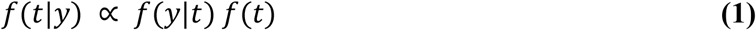

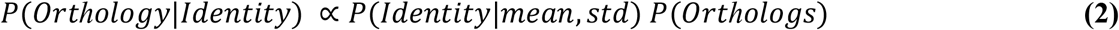

where *y* is a vector of protein sequence identities corresponding to the best hits of an alien in each of the proteomes included in the phylogenetic shadow, and *t* represents the vertical inheritance model derived from comparing the full proteomes included in the phylogenetic shadow in a pairwise fashion. The distribution *f*(*y*|*t*) is the sampling density for *y*, describing the probability that an ‘alien’ gene with an identity higher than 55% follows the vertical inheritance model; *f*(*t*) is the prior phylogenetic distribution determined by the probability of orthologs (*P*_*0*_) between the query species and the species in the phylogenetic shadow; finally *f*(*t*|*y*) is the posterior distribution of *t*. Log-likelihoods for all alien genes detected by the parametric component in a given recipient genome were derived from Eq 2. To determine the number of orthologs between the species, we used orthoMCL-pipeline (https://github.com/apetkau/orthomcl-pipeline). To derive the vertical inheritance model (*P*(*Identity*|*mean, std*)), some considerations were taken like (*i*) the mean and standard deviation of all protein sequence identities resulted from the global alignments between each proteome pair, (*ii*) we only considered putative alien genes longer than 70 amino acids. The log-likelihoods of all alien genes were classified in two classes with fuzzy clustering (Fig 1C). The group with lower values of likelihood represented the genes predicted as HGT events and the other the vertically inherited genes with an unusual composition.

As shown, our phylogenetic inference is based on a phylogenetic shadowing model instead of deriving a phylogenetic tree. To construct this model, it is important to have enough species that are closely-related to the query sequence in order to detect homology, but also a correct number of species that explains the divergence gradually. The user has to specify a priori a set of proteomes of different species to construct the phylogenetic shadow. For more information about how the species are selected, see S1 Text.

ShadowCaster algorithm was mainly implemented in Python but shares functions of R and Perl too. There are minimal external dependencies (*e.g.* OrthoMCL [22], Blastp and EMBOSS package for running the phylogenetic component). A script to build the phylogenetic shadow is provided to help the user with the input requirements (more information is supplied in the documentation of ShadowCaster).

## Results

### Performance on simulated data

To determine the robustness of ShadowCaster’s approach, we conducted experiments using simulated sets from artificial gene transfers of different origins. For this purpose, an artificial genome was modeled to be the recipient sequence based on *Escherichia coli K-12 substr. MG1655* genome. Native genes were extracted from the genome using k-mers and codon usage properties, this step ensures the removal of highly atypical genes. We created three data sets with transfers from different phylogenetic distance species like *Methanocaldococcus jannaschii* (far), *Sinorhizobium meliloti* (medium-far) and *Salmonella enterica* (close). For each set, we randomly transferred ten genes that had no orthology with the recipient sequence from the donor species previously mentioned.

We tested the effect of two parameters of ShadowCaster: *nu* parameter and the number of proteomes needed to construct the phylogenetic shadow. The *nu* parameter used in the parametric component was varied from 0.1 to 1.0 and the number of proteomes was changed from 10 to 30. The purpose of these tests is to find the optimal combination of these parameters to achieve the best performance of ShadowCaster depending on the origin of HGT events.

Fig 2 shows the curves for True and False Positive Rates of the tested parameters. The influence of the *nu* parameter was only evaluated on the HGT detection within the group alien/atypical genes. The following conclusions can be made from these curves:
1. ShadowCaster identifies with the highest precision the HGT events from medium-far and far donors due to the differences in nucleotide composition content and orthologs.
2. At the end of the parametric component, close HGT events are complex to classify within the group of atypical/alien genes due to the similarity they share with the recipient sequence. This issue can be solved by incrementing the *nu* value but it is important to emphasize that will also increase the false positive rate too.
3. A phylogenetic shadow built with 25 proteomes provides enough information to identify HGT events with the lowest probability of misclassification regardless of the origin of the event.

**Fig 2.**
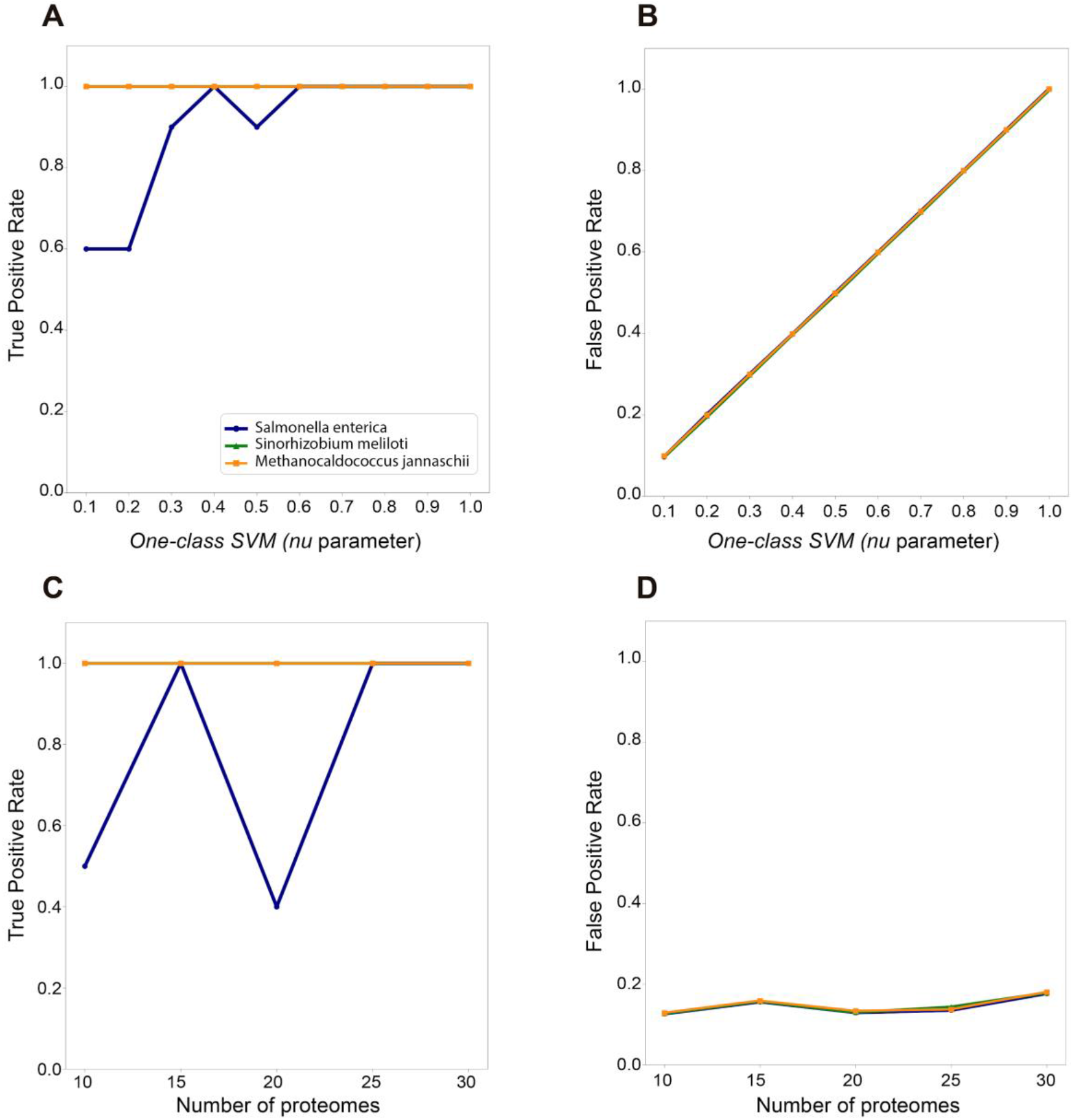
True positive and False positive rate curves for showing the performance of the two user defined parameters (*nu* and number of proteomes) of ShadowCaster.

### Performance on a real dataset. Comparison with the most popular state-of-the-art computational tools

To show the purpose of using a hybrid approach in the detection of HGT events, we validate ShadowCaster with a real genome, *Rhodanobacter denitrificans* 2APBS1, retrieved from Hemme et al. [3]. In their work, they analyzed fifty-one genes related to heavy metal resistance in the metagenome of *Rhodanobacter* populations living in contaminated-groundwater. A total of 39 genes were identified as HGT genes by at least one bioinformatic tool. We compare our results to the two best performing tools applied on their work, DarkHorse and AlienHunter, and also to another of the state-of-the art methods based on blast searches, HGTector.

As we ignore the origin of the HGT events, we took the best values obtained from the experiments with the three simulated datasets to run ShadowCaster, the *nu* parameter was set to 0.4 and the number of proteomes to build the phylogenetic shadow to 25. With the aim of comparison, DarkHorse and AlienHunter were run in the same way as mentioned in Hemme’s methods [3]. For more details about the parameter settings used to run the three tools, see Table S1. The results of ShadowCaster and the other methods are summarized in Fig 3. The use of our hybrid approach allows identifying a consensus of genes that are separately identified by the other methods. ShadowCaster and AlienHunter were the two best tools to detect HGT genes associated with metal resistance.

**Fig 3.**
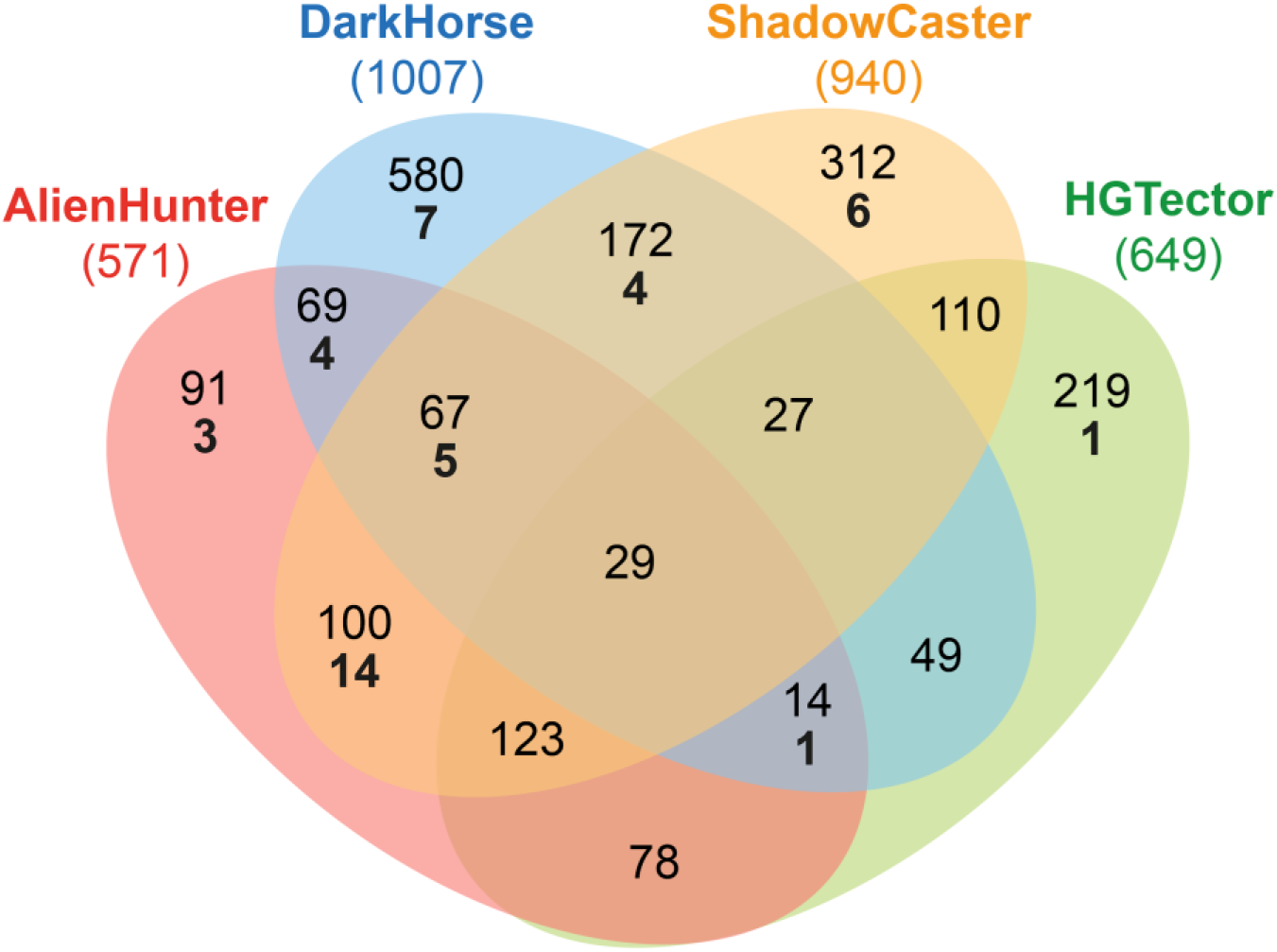
Venn diagram illustrating the HGT events detected by three of the state-of-the-art computational tools (Alien Hunter, DarkHorse and HGTector), and by the presented methodology ShadowCaster in the genome of *Rhodanobacter denitrificans* 2APBS1. HGT predictions performed by each tool is framed inside a coloured ellipse. All HGT detections are shown for each tool (black numbers inside each ellipse): Alien Hunter (571), DarkHorse (1007), HGTector (649) and ShadowCaster (940) while HGT events related to heavy metal resistance are labelled in bold numbers: Alien Hunter (27), DarkHorse (21), HGTector (2) and ShadowCaster (29).

In prokaryotes, genes acquired horizontally typically tend to code enzymes that are responsible for the growth of metabolic networks [23]. Because this dataset represents a microbial population with extreme exposure to heavy metal contamination, we wanted to know the contribution degree of the genes detected as HGT to the metabolic pathways for heavy metals. The analysis of Gene Ontology (GO) terms (Fig 4) shows that the genes predicted by ShadowCaster enrich more for enzymatic activities/systems related to heavy metals metabolism than those of AlienHunter. These results confirm the benefits to use a hybrid approach in the detection of HGT events.

**Fig 4.**
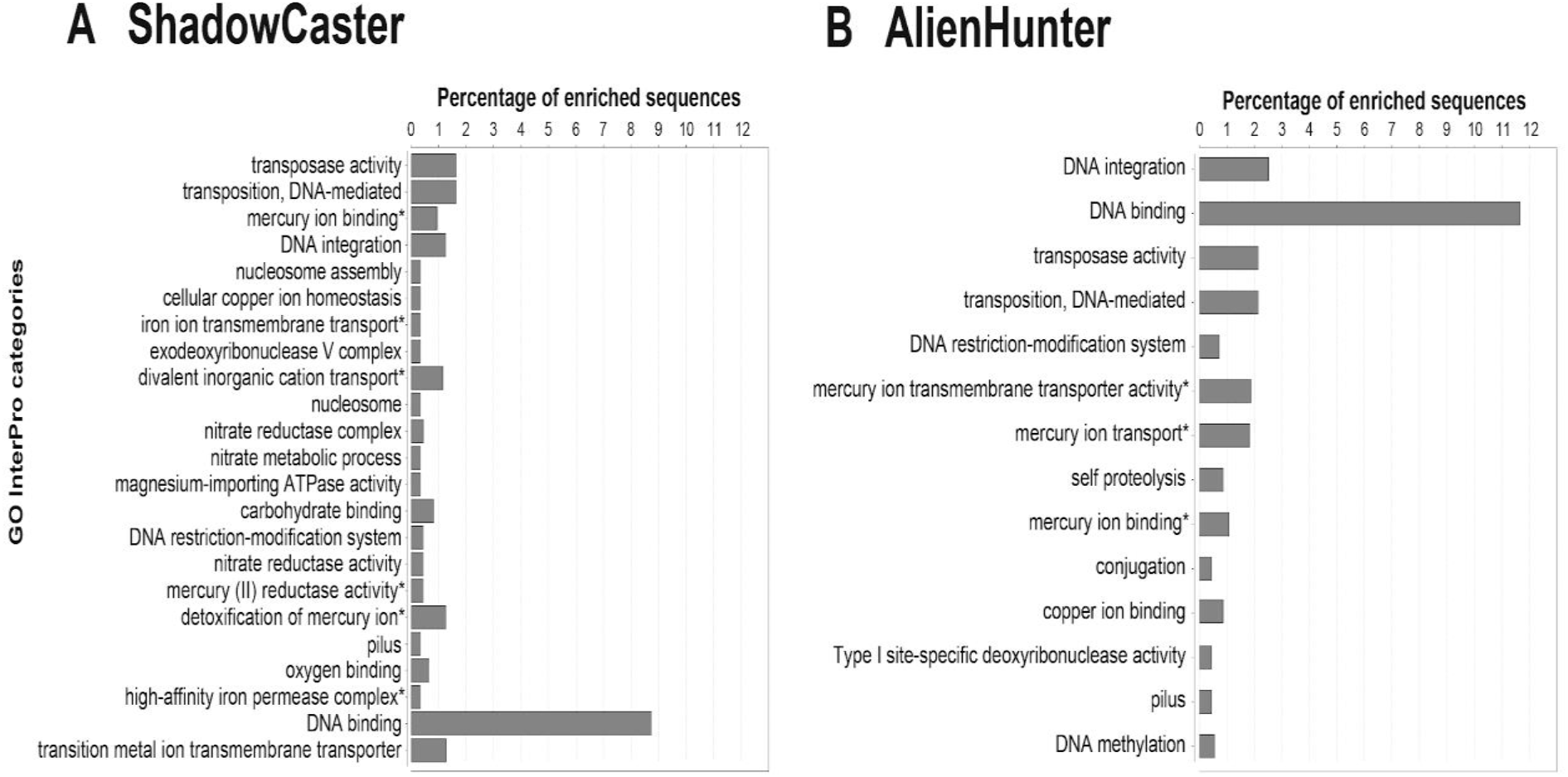
Gene Ontology (GO) Enrichment Analysis. Distribution of GO-InterPro terms exhibiting statistical significance difference (Fisher Exact Test, filtering p-values for multiple testing using False Discovery Rate) for all HGT detections performed by ShadowCaster (A) and Alien Hunter (B) in the genome of *Rhodanobacter denitrificans* 2APBS1. The analysis was conducted using the Blast2GO PRO version.

## Availability and future directions

ShadowCaster is released as an open-source software under the GPLv3 license. Source code is hosted at https://github.com/dani2s/ShadowCaster and documentation at https://shadowcaster.readthedocs.io/en/latest/. Data used for the analysis of *R. denitrificans* with an example of ShadowCaster’s output is available at https://github.com/dani2s/ShadowCaster_testData.

In future versions, we intend to detect HGT in metagenomes. Thus, several additional functions will be added to ShadowCaster such as (i) select a potential acceptor genome within a metagenome (ii) detect the HGT that most probably occurred within the metagenome based on mining the sequence of the selected acceptor and its shadow (iii) map prioritized HGT within the metagenome (acceptor and donor) (iv) iterate the search across all potential acceptors (v) draw the HGT network i.e. acceptor-donor pairs for the entire metagenome.

## Supporting Information

**S1 Text.** Proteomes selection for constructing a phylogenetic shadow

**S1 Text (Table S1).** Settings used for running state-of-art methods and output processing during the detection of HGT events in *R. denitrificans 2APBS1.*

## Acknowledgments

GACh was supported through national funds provided by the FCT - Fundação para a Ciência e a Tecnologia (FCT), I.P (UID/Multi/04423/2019).

