## Supplementary material for "ShadowCaster: compositional methods under the shadow of phylogenetic models for the detection of horizontal gene transfer events in prokaryotes": S1 Text

### **Supporting information**

**S1 Text. Proteomes selection for constructing a phylogenetic shadow**

In this section, we describe the criteria to select the proteomes to construct the phylogenetic shadow. As you can see in the documentation of ShadowCaster, the user can provide the list of proteomes by two ways: using the helper script that download the proteomes from the NCBI ftp site or a collection of proteomes (fasta file) selected by the user.

**The weight of each taxonomic rank in the phylogenetic shadow**

The phylogenetic shadow is constructed by species from different phylogenetic distances based on the taxonomy of the query species. In order to describe the divergence, our phylogenetic shadowing model is built with proteomes from nearly related species. The proteomes retrieved by the helper script are distributed according to the following taxonomic weights related with the query species: 40% from the family, 20% order, 20% phylum, 12% kingdom and 8% of another kingdom that belongs to prokaryotes. In case that there are not enough sequences to complete the percentage assigned to any of the ranks, the difference will be added to the upper taxonomic rank in order to complete with the total number of sequences to construct the shadow.

An example, if we consider *Escherichia coli* as the query species. Its phylogenetic shadow will be composed by unique protein sequences of species of the following taxonomic ranks: 40% of proteomes from *Enterobacteriaceae,* 20% *Enterobacterales,* 20% *Proteobacteria,* 12% *Bacteria,* and 8% *Archaea.*

### **Table S1. Settings used for running state-of-art methods and output processing during the detection of HGT events in *R. denitrificans 2APBS1.***

| **Method** | **Input parameters** | **Output processing** |
| --- | --- | --- |
| AlienHunter | -c optimize predicted boundaries with a change-point detection 2 state 2nd order HMM | Regions with score => 20 |
| DarkHorse | Blast_search_(all_*R.denitrificans 2APBS1*_proteins_*vs*_reference database)_table_filtered_query_coverage=70  BLAST Hit Filtering:  -filter_threshold= 0.1  -min_lineage_terms= 3  -min_align_coverage= 0.7  -Excluding terms:  Rhodanobacter  clone  cloning  construct  contaminant  cosmid  expression  plasmid  synthetic  vector | Lineage probability index(LPI) <= 0.8 |
| HGTector | BLAST parameters: evalue=1e-20  -identity=30, -coverage=50  Hit Filtering:  -ignore Taxa=unknown, uncultured, unidentified, unclassified, plasmid, vector, synthetic, phage, sp. 2APBS1  Algorithm parameters:  -howCO=4  -globalCO=0.25  -exOutlier=3  -dipTest=1  -dipSig=0.05  -toolKDE=1  -bwF=0.3  -toolExtrema=1 | modKCO=1  Conservative cutoff (default): the midpoint (arithmetic mean) of the x-coordinates of the first peak and the first pit |
